# Firing patterns of ventral hippocampal neurons predict the exploration of anxiogenic locations

**DOI:** 10.1101/2022.03.22.485343

**Authors:** Hugo Malagon-Vina, Stéphane Ciocchi, Thomas Klausberger

## Abstract

The ventral hippocampus (vH) plays a crucial role in anxiety-related behaviour and vH neurons increase their firing when animals explore anxiogenic environments. However, if and how such neuronal activity induces or restricts the exploration of an anxiogenic location remains unexplained. Here, we developed a novel behavioural paradigm to motivate rats to explore an anxiogenic area. Rats ran along an elevated linear maze with protective sidewalls, which were subsequently removed in parts of the track to introduce an anxiogenic location. We recorded neuronal action potentials during task performance and found that vH neurons exhibited remapping of activity, overrepresenting anxiogenic locations. Direction-dependent firing was homogenised by the anxiogenic experience. We further showed that the activity of vH neurons predicted the extent of exploration of the anxiogenic location. Our data suggest that anxiety-related firing does not solely depend on the exploration of anxiogenic environments, but also on intentions to explore them.

## Introduction

In the *Epistulae Morales ad Lucilium,* Seneca wrote: “There are more things, Lucilius, likely to frighten us than there are to crush us; we suffer more often in imagination than in reality”. This sentence, from one of the key figures of the school of stoicism, describes inner fear within our imagination in the absence of a direct fear-provoking stimulus. Nowadays, even though anxiety and fear correspond to the same theoretical construct in some literature, anxiety differentiates from the latter based on the potential nature of the threat in the absence of an imminent harmful stimulus(Calhoon and Tye, 2015, Davis et al., 2010, Steimer, 2002). Anxiety disorders are becoming more commonly reported: 12-month prevalence estimates on mental disorders show that at least 14% of people in the European Union suffer from anxiety disorders(Wittchen et al., 2011), and around 31% of people in the United States have experienced some type of anxiety disorders in their lifetime(Kessler et al., 1994). Different brain areas play a role in the underlying circuitry of anxiety(Sandford et al., 2000). Stimulations in the brainstem, more precisely in the periaqueductal grey matter (PAG) or the locus coerulus are specifically involved in the symptomatology of anxiety(Graeff et al., 1993, Redmond and Huang, 1979). Some studies have also shown how the amygdala plays a role in humans suffering from anxiety disorders(Davidson et al., 1999, Birbaumer et al., 1998) or in animal models with generalised fear responses associated with anxiety(Tovote et al., 2015, Wolff et al., 2014, Grundemann et al., 2019). Also, by using the elevated plus maze (EPM) as an anxiety task(Pellow et al., 1985), Tye and colleagues induced anxiolytic effects by targeting projections from the basolateral amygdala (BA) to the central nucleus of the amygdala(Tye et al., 2011). Additional structures are involved in anxiety and include the bed nucleus of the stria terminalis, whose subdivisions are differentially involved in anxiety responses(Duvarci et al., 2009, Kim et al., 2013), and the medial prefrontal cortex (mPFC), which has also been directly linked to anxiety processing in both humans(Rauch et al., 1997) and rodents(Park et al., 2016, Shah and Treit, 2003).

Last but not least, the ventral hippocampus (vH) plays a critical role in anxiety behaviour. Lesions in the vH induce anxiolysis during the exploration of elevated open arenas (e.g. EPM)(Kjelstrup et al., 2002, Padilla-Coreano et al., 2016, Jimenez et al., 2018), or generally in tasks associated with approach-avoidance conflicts(Schumacher et al., 2018). Neurons recorded in the vH showed increased firing in locations with elevated anxiety(Jimenez et al., 2018, Ciocchi et al., 2015). Likewise, projections neurons from- and to the vH exhibit anxiety-related activity: amygdala projections to the vH are specifically shaping anxiety-related behaviour during the exploration of an EPM(Felix-Ortiz and Tye, 2014, Pi et al., 2020). Similarly, the reciprocal connection (from vH to BA) is involved in the expression of context-dependent fear memories(Kim and Cho, 2020). Furthermore, information related to anxiety in the vH is directly routed to the mPFC(Ciocchi et al., 2015), and synchronised activity in this monosynaptically connected long-range circuit(Adhikari et al., 2011, Adhikari et al., 2010) is essential for the expression of anxiety behaviour. Motivated by the critical role of the dorsal hippocampus region (dH) in the encoding of spatial information(O’Keefe, 1976, Okeefe and Nadel, 1979), several studies focusing on the vH have described its involvement in spatial coding (Poucet et al., 1994, Jung et al., 1994). Yet, probably due to its anatomical location and the difficulties to record action potentials of single-unit activity in the vH, the individual neuronal dynamics associated with the changes between anxiogenic and non-anxiogenic states during spatial navigation remain poorly understood.

To study the neuronal dynamics associated with anxiety behaviour in the vH, we simultaneously recorded the activity of individual neurons during the exploration of the EPM, as well as during the exploration of a novel behavioural paradigm, the elevated linear maze (ELM). During the same recording session, we modified the ELM from a non-anxiogenic to an anxiogenic configuration, while recording the activity of the same individual vH neurons. This enabled the investigation of the neuronal dynamics within the vH underlying different anxiety states. We specifically examined the remapping of neuronal activity at the single neuron- and population-level as animals transitioned from a non-anxiogenic configuration to an anxiogenic one. Collectively, the results of this study show that the neuronal activity in the vH does not simply reflect anxiogenic locations but that it is dynamically modulated by the experience and expectation of anxiety during spatial navigation.

## Results

### The firing activity of ventral hippocampal neurons is dynamically modulated during EPM exploration

Rats (*n* = 6) freely explored an EPM consisting of two opposite arms with protective side-walls (closed arms) and two opposite arms without sidewalls (open arms) (Figure 1A). The rats exhibited strong anxiety-related behaviour by spending most of the time in the closed arms, avoiding the more anxiogenic open arms and the centre (Figure 1A, bottom. Closed vs open, *p* = 9.5615e-10; closed vs centre, *p* = 9.5657e-10. One-way ANOVA, Tukey-Kramer for multiple comparisons). While rats explored the EPM, we recorded neuronal activity in the vH with tetrodes (Figure 1B) and isolated individual spikes from different single neurons (*n* = 98). We identified vH neurons with previously described activity patterns(Ciocchi et al., 2015), exhibiting preferential firing in the open arms, closed arms, or centre of the EPM (Figure 1C). To understand the effect of the open areas on the neuronal activity, we defined six possible trajectories taken by animals during EPM exploration (from one closed arm to any other arm). After linearizing the trajectories (see methods), we organised the activity of the recorded neurons based on the spatial location of their peak (i.e. maximal) firing activity (Figure 1D). We observed that the peak firing activities of individual neurons spanned the entire maze even though the exploration of open arms was minimal. When plotting the peak firing density across the maze (normalised by the total number of neurons possibly firing in a spatial bin) the activity of vH neurons was concentrated around the centre of the maze when shuttling from one closed arm to the other one (Figure 1E top, p(C2-C1) = 0.0038, p(C1-C2) = 0.0352; bootstrapping). In some trajectories, a similar effect is observed when animals shuttled from one closed arm to an open one (Figure 1E bottom, p(C1-O2) = 0.67, p(C1-O1) = 0.0108, p(C2-O1) = 0.79, p(C2-O2) = 0.1654; bootstrapping). Even though the reduced number of entries to the open arms and exploration of open areas prevents a powerful statistic calculation, the observed effects imply that open areas are relevant for the neuronal activity of the vH.

**Figure 1.**
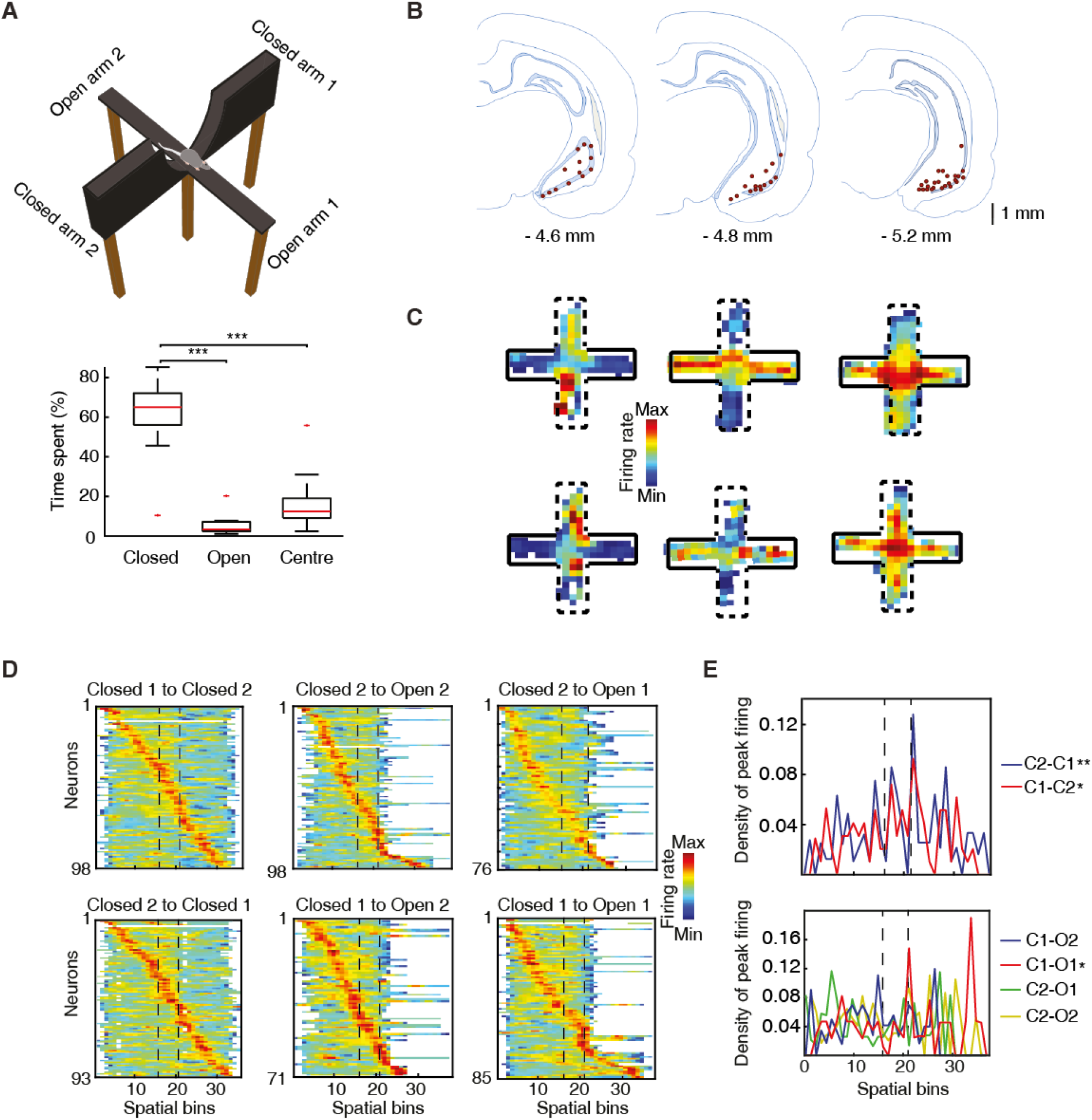
The activity of ventral hippocampal neurons is dynamically modulated during elevated-plus-maze exploration. (**A**). Top, picture of the EPM. Bottom, the percentage of the time spent in different areas by the rats (*n* = 6) during the exploration of the EPM. The time spent in closed arms is significantly higher than in the open arms and the centre (p = 9.5615e-10, p = 9.5657e-10, respectively. One-way ANOVA, Tukey-Kramer for multiple comparisons) (**B**). Location of the recording in the ventral hippocampus are indicated by red dots in three consecutive coronal sections (*n* = 47, number of rats = 8, one additional rat, for which histological location could not be confirmed, was included based on: insertion coordinates, oscillatory LFP profile and similarity of neuronal activity) (**C**). Firing rate of 6 individual neurons during the exploration of the EPM. Full black lines denote a closed arm while the dotted lines indicate an open arm. Note three different anxiety-related activity patterns: increased firing in the open arms (left) or in the closed arms (centre) or in the centre (right). (**D**). Z-transformed firing rates (colour-coded) of ventral hippocampal neurons during the exploration of the EPM, separated by trajectories and sorted by the spatial location of their peak firing activity. Dotted lines indicated the centre area. (**E**). Top, density plot of the peak firing activity location for all neurons recorded during the journeys from a closed arm to the other during EPM exploration (p(C2-C1) = 0.0038, p(C1-C2) = 0.0352, bootstrapping). Note the increased number of peak activity at the centre (i.e. the only open area for these trajectories). Bottom, same as on the top, but for the trajectories between a closed arm and an open arm (p(C1-O2) = 0.67, p(C1-O1) = 0.0108, p(C2-O1) = 0.79, p(C2-O2) = 0.1654, bootstrapping). The dotted lines denote the beginning and end of the centre area.

However, these results suffer from important limitations: First, the extremely low and sporadic exploration of the open arms does not provide robust data sampling to test hypotheses regarding the neuronal computations associated with the exploration of an anxiogenic location. Second, the non-anxiogenic and the anxiogenic areas of the EPM are constantly present. This means that the characterization of the anxiety states experienced by animals is not trivial as these may feel continuously anxious not solely in open spaces but also when considering to visit an open arm, while being in the closed arms of the EPM. In order to discriminate between neuronal activity related to spatial exploration from anxiety-related exploration and to motivate the exploration of anxiogenic areas for quantitative evaluation of the associated neuronal activity, we developed a novel behavioural paradigm to better control the transitions between anxiety states and the extent of exploration of anxiogenic areas.

### Removal of protective sidewalls along an ELM induces anxiety behaviour

We developed an ELM, which consisted of an elevated linear track with removable protective sidewalls. This allowed to have sidewalls either all along the entire ELM, called closed-closed configuration (CC), or to have the sidewalls removed from half of the track, called closed-open (CO) configuration (Figure 2A). The rapid removal of the sidewalls enabled the alternation between a non-anxiogenic and an anxiogenic configuration within the same maze and in a single recording session. To control for neuronal activity-associated differences between two dissimilar areas in the linear maze exploration, while maintaining non-anxiogenic locations, we modified temporarily the visual appearance of sidewalls and floor texture in one half of the track, creating a new and enriched environment, without being anxiogenic. This configuration was termed closed-texture (CT) configuration (Figure 2B). Rats (n = 6) were motivated to fully explore the ELM, by shuttling from one end of the track to the other one over numerous trials, to receive food rewards (Figure 2C). To assess whether the removal of sidewalls induces behavioural readouts of anxiety similar to those of the EPM, we calculated the percentage of time spent on each of the arms for each configuration (Figure 2D). No differences were observed in the time spent by rats on the different arms during non-anxiogenic explorations (CC, CT). On the contrary, a significant difference was found in the configuration with sidewalls removed (CO exploration, p = 1.45e-05, Wilcoxon signed-rank). The time spent in the centre area (11.5 cm around the middle of the track) was also longer during CO exploration compared to CC or CT explorations (Figure 2E, p = 2e-05 and p = 0.034 against CC and CT respectively, one-way ANOVA, Turkey-Kramer for multiple comparisons), suggesting of hesitations to enter this open area of the maze.

**Figure 2.**
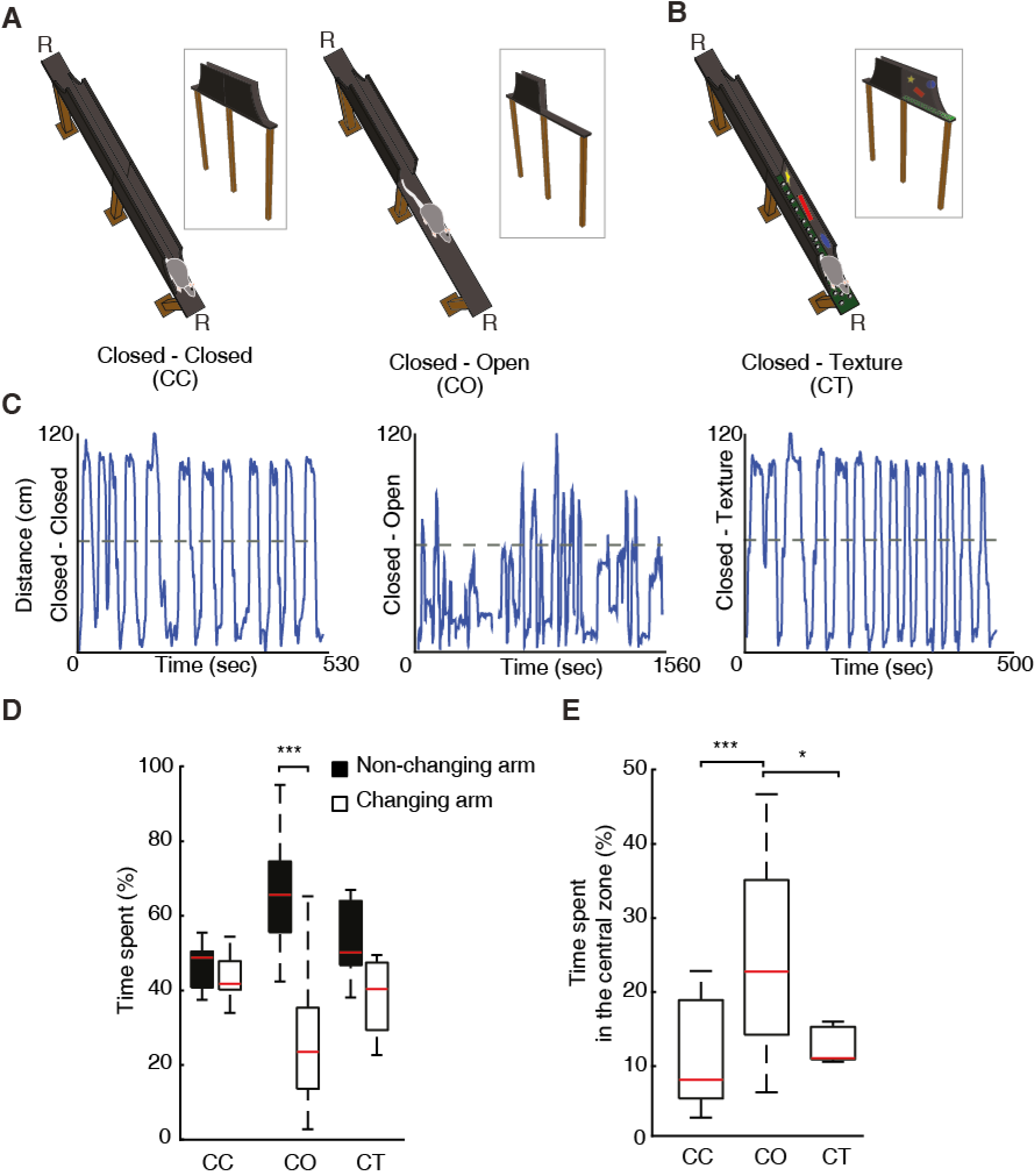
Removal of protective sidewalls along an elevated-linear-maze induces anxiety behaviour. (**A**). ELM configurations with sidewalls along the entire track (CC, both arms closed, left), and with sidewalls removed for half of the track (CO, called one arm closed – one arm open, right). Note the presence of a non-changing arm, while the other arm changes from a closed to an open configuration. R, indicates locations of food reward. (**B**). ELM configuration with both arms closed, but the floor texture and visual cues on inner-walls are changed in one of the arms (CT, closed and texture arm). (**C**). Linearised trajectories of a rat running in the three different ELM configurations during a single behavioural session. The grey line denotes the centre of the linear maze and the division between the two arms. (**D**). Time spent in both the non-changing and the changing arm in each configuration. Significant differences in the time spent appeared solely in the CO configuration (p = 1.45e-05, Wilcoxon signed-rank). (**E**). More time spent in the central zone (defined as 11.5 cm around centre of the track) in the CO configuration in comparison to the CC and CT configurations (p = 2e-05 and p = 0.034 respectively, one-way ANOVA, Tukey-Kramer for multiple comparisons).

Overall, the removal of sidewalls along the ELM induced anxiety behaviour that evolved during single behavioural sessions according to the anxiety content of each maze configuration.

### Overrepresentation and remapping of vH activity during anxiety

We recorded a total of 133 neurons with tetrodes in the vH (Figure 1B), while the rats were exposed to the different configurations of the ELM. When the activity of individual neurons was sorted according to the spatial location of their peak firing activity, we observed that it spanned over the entire extent of the ELM for the three different configurations (Figure 3A). However, during the CO exploration, the entire distribution of peak firing activity was skewed toward the open area. We assessed the proportion of vH neurons with peak firing activity located on the different halves of the ELM during CC exploration. We found no differences in the proportion of peak firing activity located in each half of the CC configuration, even for the closed half that was going to be opened in the CO configuration (Figure 3B, left). Then, when one half was opened, a remapping of the neuronal activity was induced towards the anxiogenic location. We observed a higher proportion of peak firing activities located on the open arm (Figure 3B left, p = 0.0121, chi-square test). In a subsequent analysis, we detected that a significant proportion of vH neurons changed the location of their peak firing activities from the closed area to the open area after sidewall removal (Figure 3B right, p = 0.0066, chi-square test). Figure 3D shows the peak activity transition of every single recorded neuron before (CC) and after (CO) removing the sidewalls in half of the maze. Importantly, no differences were found for the location and location changes of peak firing activity during the exploration of the CT configuration using a novel texture and visual cues in the closed arm (Figure 3C), suggesting that dynamic remapping of vH activity is contingent to the experience of anxiety rather than to a stimulus-enriched environment or novelty.

**Figure 3.**
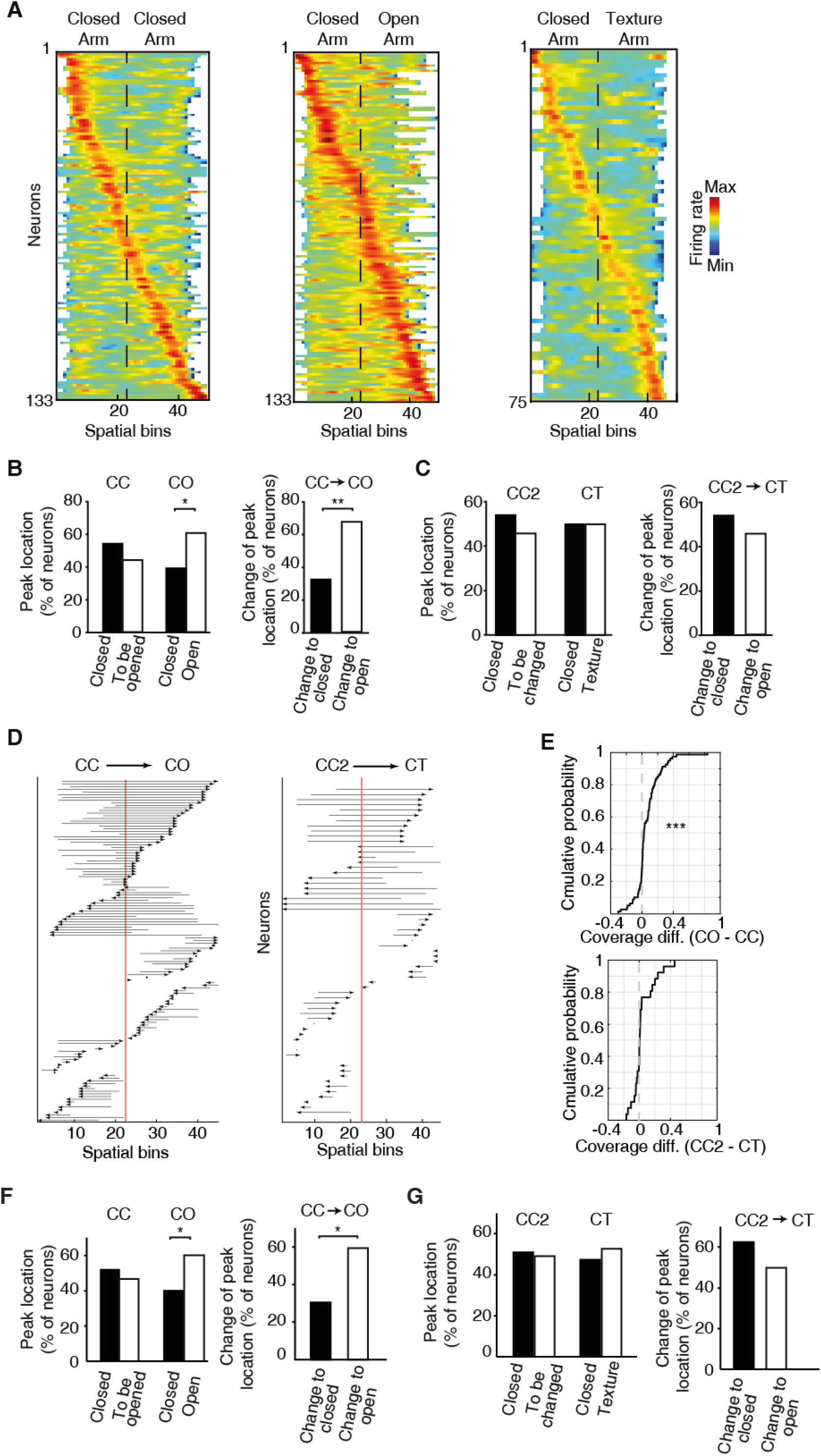
Overrepresentation and remapping of ventral hippocampal activity during anxiety. (**A**). Z-transformed firing rates (colour coded) of ventral hippocampal neurons during the exploration of the ELM and sorted by the spatial location of their peak firing activity for the three configurations: CC (left); CO (centre); CT (right). The order of neurons is sorted for each configuration independently. Dotted lines indicated the centre of the maze. Note the increased number of neurons with peak firing activity in the open arm. (**B**). Left, comparisons of the percentage of neurons with peak firing activity located in the different arms of the CC and CO configurations of ELM. Note the significant differences of the proportion of neurons with peak firing activity in the open arm (*p* = 0.0121, chi-squared test). Right, upon removal of the sidewalls, a larger proportion of neurons change the location of their peak firing activity from a previously closed to a currently opened arm (*p* = 0.0066, chi-square test). (**C**). Same analysis as in (B) for the CT configuration. CC2 is a configuration with sidewalls along the entire track (fully closed) explored right before the presentation of the CT configuration. No significant differences were found (**D**). Changes of the spatial location of the peak firing activity for individual neurons between different configurations. Each arrow denotes the remapping of the location of the peak firing activity between the CC (base of the arrow) and the CO configurations (arrowhead). The red line indicates the centre of the linear maze. Note that the peak activity of the neurons shifted towards the open arm when changing from the CC to CO configuration. (**E**). Comparison between the coverage index of the activity of single neurons (see methods) with peak firing located in the opened arm, recorded during ELM exploration. Top, the cumulative probability of the distribution generated by subtracting the coverage indexes of the CC configuration from the coverage indexes of the CO configuration. The plot indicates that the firing rate’s coverage index of individual neurons significantly increased when the rat explored the CO configuration in comparison with the CC configuration (p = 6.865e-05, Wilcoxon signed-rank). Bottom, same as left but without differences between the CC2 and CT configurations (p = 0.4237, Wilcoxon signed-rank). (**F**). Left, comparisons of the percentage of neurons with peak firing activity (after correction of the speed influence in the spiking activity by using the residuals of the GLM model) located in the different arms of the CC and CO configurations of ELM. (*p* = 0.019, chi-squared test). Right, upon removal of the sidewalls, a larger proportion of neurons change the location of their peak firing activity (after correction of the speed influence in the spiking activity) from a previously closed to a currently opened arm (*p* = 0.00176, chi-square test). (**G**). Same analysis as in (C) for the CT configuration. CC2 is a configuration with sidewalls along the entire track (fully closed) explored right before the presentation of the CT configuration. No significant differences were found.

Movement changes, mostly related to the running speed of the animal, could have been responsible for the neuronal activity changes observed during the exploration of the open arm. Speed related modulation of hippocampal activity has been widely shown in the dorsal hippocampal region(Wiener et al., 1989, McNaughton et al., 1983, Czurko et al., 1999). Using generalized linear models (GLMs), we aimed to capture the influence of the animal’s instantaneous speed on the location dependent activity for each cell (see methods). We found that the spiking activity of 49 neurons out of 133 (36.84%) was significantly modulated by the speed of the animal during the exploration of the CC configuration. Also, as expected, a higher number of neurons (72 out of 133, 54.14%) were significantly modulated by running speed during the exploration of the CO, in agreement with the speed changes related to the experience of an anxiogenic area. Next, we used the residuals of the GLM model as an approximation of the neuronal spike-associated activity, corrected by the influence of the speed. Repeating the same analysis as in Figure 3B,C we found that a significant proportion of vH neurons changed the location of their peak firing activities from the closed area to the open area after sidewall removal (Figure 3F), in addition to no significant differences when exploring the CT configuration (Figure 3G). These results suggest that even though there is a modulation by the speed of the animal, there is also a prominent influence of the anxiogenic location in the neuronal activity irrespective of running speed.

Additionally, we observed (Figure 3A) that the activity of vH neurons appeared to be spatially broader during CO exploration, suggesting that the anxiogenic area in the CO configuration selectively affect the spatial features of vH neurons. To validate the spatial broadening of the neuronal activity in the CO configuration, we used a coverage index (also known in the literature as sparsity index(Skaggs et al., 1996), see methods). The index measures the spatial coverage of the activity of a neuron. Values close to 1 indicate that the activity of the neuron covers almost the entire linear maze (e.g. a value of 0.9 implies a coverage of 90% of the space). On the contrary, values close to zero imply that the activity of the neuron is spatially selective. We compared the change of this index during changes in ELM configuration (CC to CO and to CT). We observed an increased coverage of neuronal activity from the CC to the CO configuration (Figure 3E top, n = 81, p = 6.865e-05, Wilcoxon signed-rank) in neurons with their main activity located in the open area. In contrast, the coverage did not change significantly during the transition between the CC and the CT configuration (Figure 3E bottom, n = 38, p = 0.4237, Wilcoxon signed-rank) in neurons with their main activity located in the new texture area.

Collectively, these data indicate a dynamical recruitment of vH neurons when swapping between a non-anxiogenic (CC) to an anxiogenic configuration (CO), generating not solely a remapping of activity but also an overrepresentation of the anxiogenic area. As these effects were not detected during a novel, but non-anxiogenic experience, we concluded that they could be attributed to the processing of anxiety during open arm exploration rather than to a changed environment or novelty perception *per se.*

### The direction-dependent activity of vH neurons becomes homogenised following the introduction of an anxiogenic location

Hippocampal place cells have been reported to exhibit direction-specific spatial modulation of activity as animals run along a linear maze(McNaughton et al., 1983, Muller et al., 1994, Royer et al., 2010).

This raises the question as to whether the activity of vH neurons, recruited by areas with increased anxiety content, are also modulated by the direction of the journey along a linear track. When monitoring the activity of individual vH neurons recorded during the exploration of the CC configuration (Figure 4A), we expectedly observed a profound direction-dependent difference. However, this direction-dependent neuronal firing of the same vH neurons became homogenised, meaning that it was very similar in both directions, once the sidewalls were removed and the rats were exploring the CO configuration. To quantify this phenomenon, we use the spearman correlation as a measurement of place field similarity (PFS) of a neuron between its activity while the animal moved in one direction vs. the other direction. The PFS value obtained from the correlation will grant an accurate approximation of the similarity between the neuronal activity of both directions. As expected, the PFS index of animals exploring the CC configuration was different from the PFS index of animals exploring the CO configuration (Figure 4B, left, p = 1.536e-08, two-sample Kolmogorov-Smirnov test). Based on the values of the PFS indices (median PFS_CC_ = −0.0539, median PFS_CO_ = 0.5091), we inferred that the significant difference between the PFS distributions is due to a similar neuronal activity in both directions during the exploration of the CO configuration.

**Figure 4.**
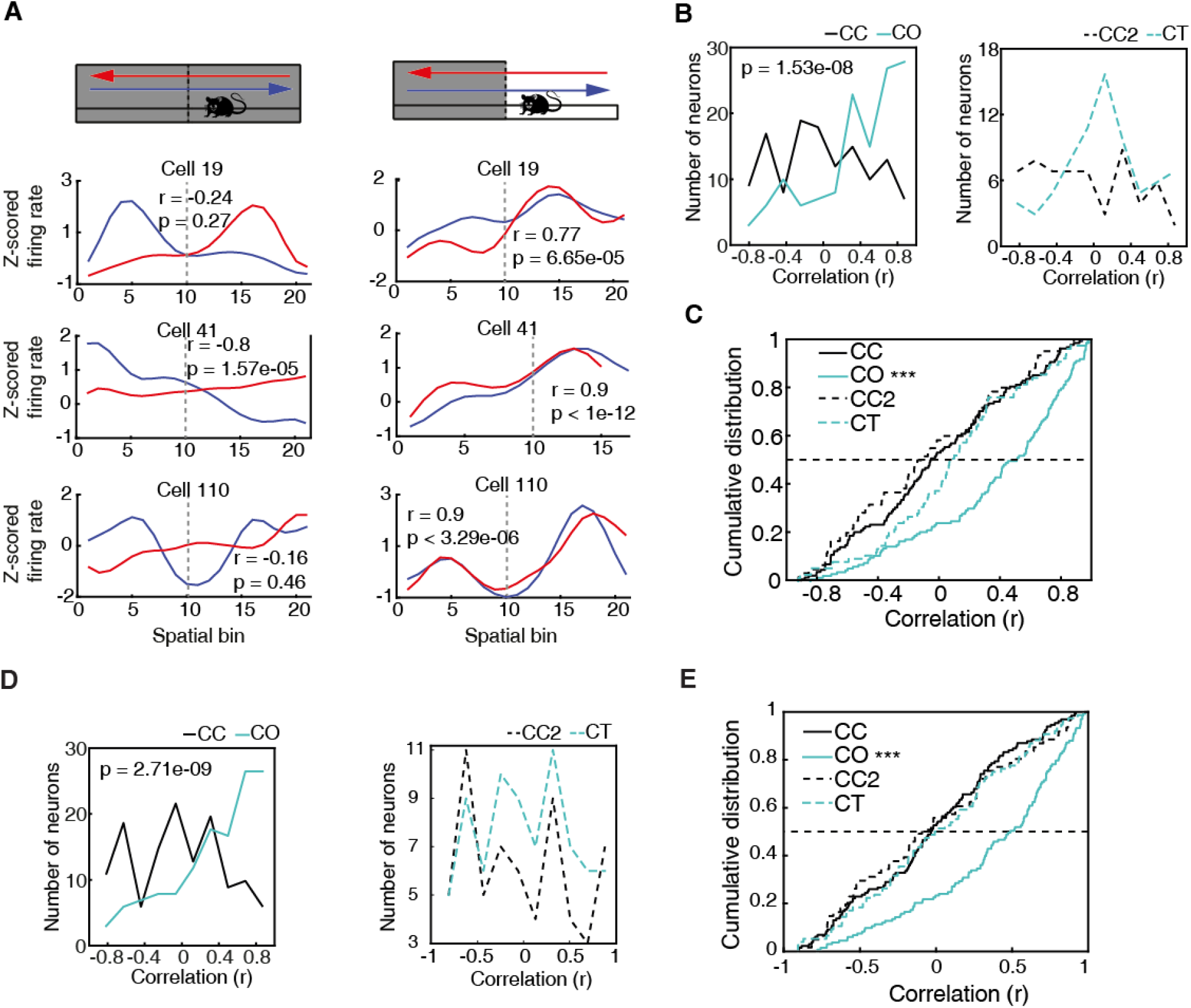
The direction-dependent activity of vCA1 neurons is homogenised in an anxiogenic location. (**A**). Neuronal activity of individual vH neurons while animals explored the ELM in both the CC and the CO configuration. Blue lines denote when animals headed towards the arm that will be open (in the case of the CC configuration) or is open (in the case of the CO configuration). Red lines denote when animals returned from this arm. Correlation values (Spearman correlation) indicate the similarity between the neuronal activities of both trajectories. (**B**). Histograms of the firing rate maps similarity index (PFS, see methods) for the activities of single neurons calculated in the two possible directions on the ELM (left to right and right to left). Left, PFS index is higher during the CO configuration (cyan) compared to the CC (black) configuration (p = 1.536e-08, two-sample Kolmogorov-Smirnov test). Right, PFS index is not significantly different during the CT configuration (dotted cyan) compared to the CC2 (dotted black) configuration (p = 0.0678, two sample Kolmogorov-Smirnov test) (**C**). Cumulative distributions of the PFS indexes for the CC, CO, CT and CC2 configurations. Note that during the CO configuration the PFS index of neurons is significantly higher compared to the other configurations (vs CC, p = 1.536e-08; vs CT, p = 8.1919e-07 and vs CCT2 p = 3.8183e-06; two-sample Kolmogorov-Smirnov test). (**D**). Histograms of the firing rate maps similarity index (PFS, see methods) for the activities of single neurons (after correction of the speed influence in the spiking activity) calculated in the two possible directions on the ELM (left to right and right to left). Left, PFS index is higher during the CO configuration (cyan) compared to the CC (black) configuration (p = 2.7102e-09, two-sample Kolmogorov-Smirnov test). Right, PFS index is not significantly different during the CT configuration (dotted cyan) compared to the CC2 dotted black) configuration (p = 0.7448, two sample Kolmogorov-Smirnov test) (**E**). Cumulative distribution of the PFS indexes in D) for the CC, CO, CT and CC2 configurations. Note that during the CO configuration the PFS index of neurons is significantly higher compared to the other configurations (vs CC, p = 2,7102e-09; vs CT, p = 3.1228e-06 and vs CC2 p = 9.1521e-06; two-sample Kolmogorov-Smirnov test).

To corroborate that such a phenomenon was not due to the novelty of the arm, we calculated the PFS indexes of the neuronal activity during the exploration of a novel arm (CT), instead of an anxiogenic location (i.e. CO configuration). The PFS indexes between the exploration of the CC configuration, prior to the CT (called CC2) and the PFS indexes during the CT configuration were not significantly different (Figure 4B, right, p = 0.0678, median PFS_CC2_ = −0.1279, median PFS_CT_ = 0.0922). In general, all the PFS values during the exploration of CC, CC2 and CT were significantly lower than during the exploration of the CO configuration, as seen in the cumulative distribution plots of the PFS (Figure 4C, CO vs CC, p = 1.536e-08; vs CT, p = 8.1919e-07 and vs CC2, p = 3.8183e-06; two-sample Kolmogorov-Smirnov test). The same tendencies were seen after obtaining the neuronal spike-associated activity controlled by the speed of the animal (Figure 4D,E), as described previously and in the methods.

In conclusion, even though there was direction-dependent neuronal activity in the vH during the exploration of a non-anxiogenic linear maze, this spatial dependency was reduced when animals encountered an anxiogenic location, and the spiking activity of the neurons tended to be homogenised independently of the direction of the animal.

### The activity of vH neurons predicts the extent of exploration of an anxiogenic location

We have shown that neurons of the vH change their activity patterns during the experience of anxiety. The described changes on neuronal activity were mainly related to the exploration of the open area. Next, we asked if neuronal activity during the exploration of the closed arm during a CO configuration, is influenced by the upcoming anxiety states. Observation of behavioural readouts (Figure 2) seem to indicate a hesitation to enter the opened arm in the CO configuration, and after entering, there is not always a full commitment to explore it to the end. Therefore, we asked whether neuronal firing in the closed area might be predictive of the extent of an upcoming exploration in the anxiogenic area. To do so, we divided spatial explorations into two groups depending on how far animals explored the open arm during a particular trial: proximal and distal exploration trials were defined using an individual threshold for each session 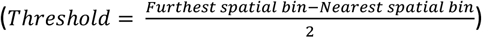 (Figure 5A). We then tested if the neuronal activity before entering the open arm was informative of how far animals would venture into the open arm (proximal or distal explorations). By using the firing activity of each vH neuron prior to the entry to the open arm in the CO configuration, we represented each trial by a population vector built with the activity of all the co-recorded neurons during single recording days by *T_n_* = 〈*FR*_1*n*_, *FR*_2*n*,_, …, *FR_mn_*〉 where FR is the firing rate calculated in the closed area, n is the total number of trials, and m is the total number of neurons. If some anxiety-related information is computed during the exploration of the closed part of the ELM, we then anticipated that this neuronal activity would predict the extent of explorations (proximal or distal explorations) in the open anxiogenic arm. To better visualise the population activity in the close arm related to the future extent of exploration of the open arm, we calculated the principal components (PCA, see methods) of the population vectors and plotted the two first principal components (Figure 5A). We observed a clear separation between the two types of trials (proximal exploration, red; distal exploration, blue) based on the neuronal activity recorded. To further strengthen this observation, we trained a support vector machine (SVM, Figure 5B,C) using the entire neuronal activity (not the PCA values) in the closed arms for each recording day and we were able to predict the extent of exploration for single trials (see methods). A set of the best predictive neurons within a session was then used for each session SVM (see methods for description on how the neurons were selected). We found that during individual recording sessions (Figure 5C), the performance of the SVM was above chance levels in 10 out of 12 sessions and approximately 24% of the neurons (32 out of 133) were predictive of proximal vs distal explorations of the opened arm by using their firing rate during the closed arm of the CO configuration.

**Figure 5.**
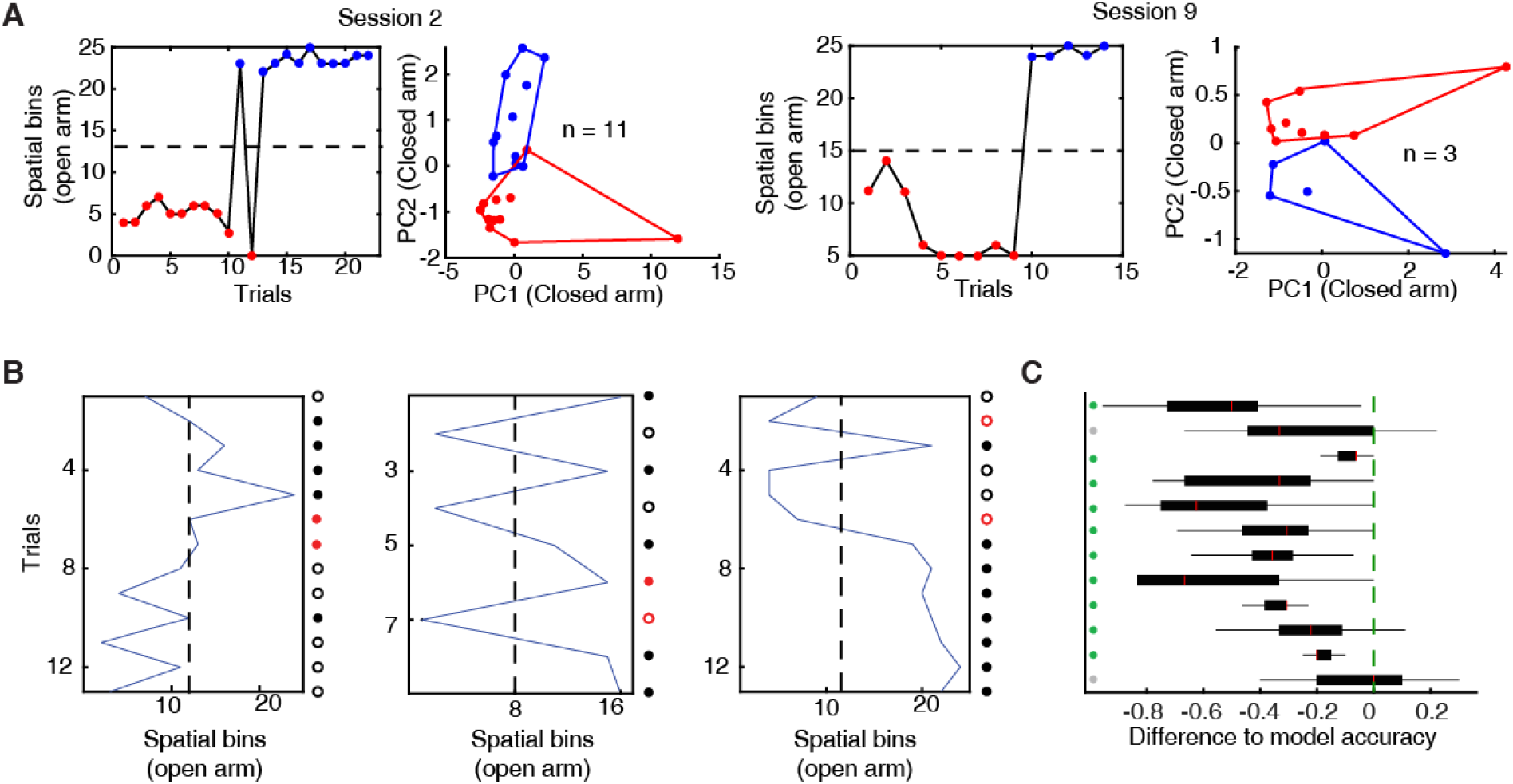
The activity of ventral hippocampal neurons predicts the extent of exploration of an anxiogenic location. (**A**). For two behavioural sessions, the furthest spatial bin visited on each trial (left) and the single-unit activity of all co-recorded neurons during the run on the closed arm projected onto their two highest principal components (right) are shown. The dotted line indicates the spatial bin set as a criterion to define two different types of trajectories: proximal (red dots) and distal (blue dots) exploration. (**B**). Predictions for three individual sessions. The blue line shows the furthest spatial bin reached during specific trials. The dotted line indicates the middle of the exploration of the open arm and divides the trials into proximal (left) and distal (right) explorations (used for the SVM-classifier). Dots at the right of each plot show the trial by trial accuracy of the SVM-classifier (see methods). Full dots show trials with distal explorations while circles show trials with proximal explorations. Red colour implies inaccurate prediction of the SVM classifier. (**C**). Normalised distributions of the SVM-classifier performance (observed data - shuffled data) for each of the 12 sessions used. The green line marks the SVM-classifier performance on the observed data. Green dots show sessions for which the SVM-classifier successfully predicted proximal or distal exploration (higher than the 95% percentile of the shuffled data) based on neuronal firing activity during the exploration of the closed arms, while grey dots show sessions with inaccurate prediction. Box plots show median, 25^th^ and 75^th^ percentile.

These analyses indicate that the neuronal representation of the anxiogenic location was not only dynamically modulated by the direct experience of anxiety, but already existed in the closed arm, possibly reflecting the intention to venture into the open arm. This implies that neuronal activity within the vH can predict upcoming anxiogenic situations, even when animals are still located in a safer environment without a direct exposure to an anxiety-inducing location.

## Discussion

To investigate the neuronal dynamics governing anxiety behaviour in the vH, we recorded the activity of individual neurons while rats explored anxiogenic locations. In addition to the classical EPM, we used a novel ELM, which allowed us to rapidly change the anxiety content of the maze to expose rats to non-anxiogenic or anxiogenic configurations. We found that the neuronal activity of the vH exhibited a uniform spatial representation in the non-anxiogenic configuration of the ELM and that vH neurons displayed direction-dependent spatial firing when shuttling from one end of the ELM to another. When the anxiogenic location was introduced by removing the protective sidewalls from half of the track, the peak firing activity remapped towards the newly introduced anxiogenic location and direction-dependent firing was homogenised. Of important note, neuronal activity in the closed arm of the ELM predicted the extent of the upcoming exploration in the open arm even before rats entered into the open anxiogenic location.

Much of the anxiety research in freely-moving rodents has been relying on the EPM. Using the EPM, it has been shown that: amygdala projections to the vH control the expression of anxiety(Felix-Ortiz and Tye, 2014, Pi et al., 2020); there is an anxiety-associated neuronal activity in the vH routed to the mPFC(Ciocchi et al., 2015); and that the vH-prefrontal pathway is critical for anxiety behaviour(Ciocchi et al., 2015, Adhikari et al., 2011, Adhikari et al., 2010). The first part of this manuscript focuses on the activity of neurons recorded in the vH during the exploration of an EPM. We divided the exploration of the EPM into different trajectories. We observed a localised increase in the density of peak firing activity when rats crossed the centre in all the different C-C or C-O trajectories (Figure 1E). This might be explained by the fact that not only the open arms of an EPM are anxiogenic, but also the centre zone(Mendes-Gomes et al., 2011). Unfortunately, it proved difficult to analyse these effects quantitatively, because rats explored the open arms only minimally and sporadically, consistent with the anxiogenic nature of the EPM paradigm.

To overcome this problem, we introduced a novel ELM on which rats were motivated to shuttle from one extremity to the other to receive rewards. The ELM had three different configurations: non-anxiogenic (CC configuration); anxiogenic (CO configuration); and a configuration with new texture and visual cues (CT). Configurations could be quickly switched within a session. At the behavioural level, we observed anxiety-related behaviour during ELM exploration comparable to the ones during EPM exploration (Figure 2D, E). However, the main advantage of the ELM was the possibility to record the neuronal activity of the same vH neurons while rapidly modifying ELM configurations and motivating rats to spend more time in the anxiogenic location. After animals transited from a non-anxiogenic to an anxiogenic configuration, we observed a significant increment in peak firing activities predominately located in the open area. We attributed this recruitment or remapping of the neuronal activity to the anxiogenic location. The neuronal mechanisms during spatial remapping remain largely elusive. Global remapping is a phenomenon observed in the dH when animals move from one environment to a different one(Leutgeb et al., 2005). One could argue that the simple removal of the walls is changing the environment of the ELM and therefore animals could perceive the open arm as a completely novel environment, inducing global remapping. Nevertheless, remapping of neuronal activity in a new environment is expected to be random and independent of a previously explored environment(Schlesiger et al., 2018, Leutgeb et al., 2005, Gauthier and Tank, 2018), even when emotional contexts are introduced(Moita et al., 2004). However, the remapping of neuronal activity observed in ventral hippocampal neurons is not arbitrary as it is most prominent in the anxiogenic location, contrary to a uniformly distributed peak activity in case of a random remapping. Nonetheless, the opening of walls *per se,* independent of the anxiogenic experience, could cause changes in vH neuronal activity. Previous work has shown that novel environments elicit the activation of vH neurons to a similar extent as an aversive stimulus(Graham et al., 2021). To show that the remapping of neuronal activity relies on anxiety, we changed the ELM from a CC to CT configuration during the same recording session. In the CT configuration, the previously open location changed to a novel one with a different floor texture and visual patterns in the inner part of the walls, but with protective sidewalls kept all along the track (Figure 2B). As anticipated, these changes equally induced a remapping of neuronal activity in the vH. Yet, in this case, the remapping was distributed over the length of the maze and the novel area did not show an increased number of peak firing activity (Figure 3C). Overrepresentation of behaviourally relevant locations is not novel in the hippocampus research field. It has been previously demonstrated that hippocampal place cells fire preferentially at reward locations during goal-directed tasks(Dupret et al., 2010, Hok et al., 2007, Hollup et al., 2001). Furthermore, as the vH is strongly associated with anxiety, the anxiogenic location represents a highly salient environment for vH-dependent computations(Bannerman et al., 2014). This is supported by our finding that the anxiogenic area is overrepresented by neurons of the vH, further strengthening its role in the emotional processing of information.

Another relevant observation relates to the directional firing of vH neurons. In the non-anxiogenic configuration of the ELM, the spatial activity of single vH neurons varied depending on the direction of exploration. Similar observations have been made in both the dH(McNaughton et al., 1983) and vH(Royer et al., 2010) for linear mazes, suggesting of a common principle underlying spatial information along the dorso-ventral axis of the hippocampus. With respect to the vH, Royer et al. hypothesised that the differential activity between inbound and outbound trajectories might be caused by the reward delivered at the end of the arm implying some reward-associated value coding. In contrast to Royer et al., we placed rewards at both extremities of the ELM. Although this does not invalidate the view of Royer et al., in our study, the direction-dependent firing was homogenised in the anxiogenic configuration of the ELM, with vH neurons exhibiting similar firing independently of the direction of exploration in the open arm (Figure 4). We attribute the homogenisation of the direction-dependent firing in single vH neurons to the relevance of the anxiogenic location for vH-dependent computations. This is further supported by the results of the control experiments, in which we introduced a new, but not anxiogenic, environment to the ELM (Figure 4C) leading to fewer changes in vH neuronal activity.

Furthermore, the modulation of the neuronal activity in the anxiogenic location was not exclusively observed while the rats explored the open area. The CO configuration of the ELM contained protective walls in half of the maze, while the other half was entirely open. The neuronal population activity was a good predictor of the extent of the exploration of the open location, even when rats explored the closed arm before entering to the open location. Indeed, the neuronal activity in the closed arm was sufficient to infer whether rats would perform proximal or distal explorations of the open arm (Figure 5). Anxiety-related modulation of vH neuronal activity in both the closed and the open locations implies that not only a direct experience of anxiety enhances neuronal activity in the vH, but also its anticipation without a direct confrontation to an anxiety-inducing situation.

Overall, we provided evidence that the neuronal dynamics within the vH are subjected to the experience of anxiety. When an anxiogenic situation was encountered, vH neurons, first, over-represented this location (Figure 3); Second, their activity was tuned to the anxiogenic environment, impairing previous direction-dependent neuronal activity manifesting in the absence of anxiety (Figure 4); Third, the neuronal activity of vH neurons reflected and predicted the exploration of an anxiogenic location (Figure 5). Collectively, these results expand our view of vH function by highlighting dynamic and predictive computations during anxiety.

## MATERIALS AND METHODS

### Experimental subjects

In total, nine long Evans rats from Charles River Laboratories (male, 250–600 g), were kept in 12 h light cycles during behavioral experiments (performed during the light cycle). All experimental procedures were performed under an approved licence of the Austrian Ministry of Science and the Medical University of Vienna.

### Surgery and microdrive implantation

Using isoflurane, animals were anaesthetised (induction 4%, maintenance 1–2%; oxygen flow 2 l/min) and fixed to a stereotaxic frame. The body temperature was controlled using a heating pad. Iodine solution was applied to disinfect the surgery site and eye cream was used to protect the eyes. Local anaesthetic (xylocain^®^ 2%) was used before the incision. Saline solution was injected subcutaneously every 2 h, to avoid dehydration. Seven stainless steel screws were anchored into the skull to improve the stability of the construct, and two of the screws were placed above the cerebellum as references for the electrophysiological recordings. Next, based on the rat brain atlas(Paxinos and Watson, 2007), a craniotomy was performed above the ventral hippocampal area (from bregma: −4.8 mm anterior, 4.5 mm lateral, right hemisphere). After removal of the dura mater, an array of 12 independently movable, gold plated (100–500 kΩ) wire tetrodes (13 μm insulated tungsten wires, California Fine Wire, Grover Beach, CA) mounted in a custom-made microdrive (Miba Machine Shop, IST Austria) were implanted (Dorso-ventral: −6.5 mm). Paraffin wax was then applied around the tetrode array, and the lower part of the microdrive was cemented (Refobacin^®^ Bone Cement) to the scalp. At the end, the surgery site was sutured, and systemic analgesia (metacam^®^ 2 mg/ml, 0.5 ml/kg) was given. Animals were allowed at least 7 days of recovery time.

### Mazes description and behavior

The elevated plus-maze (EPM) consisted of two closed (with protective side walls) and two open (without sidewalls) arms. The dimensions of the arms were 9 x 50 cm, the walls in the closed arms were 40 cm high, and the EPM was elevated 70 cm above the floor. Rats were placed on the EPM facing the open arm distal to the experimenter. Sessions lasted 5 – 8 min and were done at 200 lux of room light intensity(Walf and Frye, 2007).

The elevated linear-maze (ELM) consisted of a linear track of 120 cm length and 8 cm in width. The maze was elevated by 105 cm above the ground. A reward was given at both endpoints (two 20 mg sugar pellets). Three possible configurations were presented during the EPM exploration: A Closed-Closed configuration (CC), consisting of 4 black panels acting as side walls which covered the entire length of the track and prevented the animal from experiencing the height; a Closed-Open configuration (CO) which consisted on 2 black panels acting as walls covering one half of the maze and leaving the other half completely open, resulting in an anxiogenic area; and a third configuration was called Closed-Texture (CT) which consisted on 4 black panels acting as walls covering the entire length of the maze. The difference with the CC configuration was that in half of the maze (the half which was open in the CO configuration) coloured geometrical figures were added to the sidewalls as new visual patterns and the texture of the floor was changed. The ELM sessions were composed by the presentation of the CO and CT configurations, each preceded by a CC configuration. Depending on the motivation of animals to explore, CC and CT configurations lasted 5 – 15 minutes while CO configurations lasted between 5 – 20 minutes. When a CC configuration was preceding a CT configuration, we refer to the configuration as CC2.

Tracking of the rats’ movement was monitored by triangulating the signal from three LEDs (red, blue, green) placed on the implanted headstage and recorded at 25 frames per second by an overhead video camera (Sony).

### *In vivo* electrophysiology

Either an Axona headstage (HS-132A, 2×32 channels, Axona Ltd) or Intan headstage (2 x RHD32-channel headstage) were used to pre-amplify the extracellular electric signals from the tetrodes. For the Axona headstages, output signals were amplified 1000 × via a 64-channel amplifier and then digitised continuously with a sampling rate of 24 kHz at 16-bit resolution, using a 64-channel analogue-to-digital converter computer card (Axona Ltd). For the Intan headstages, signals were acquired with the RHD32 channel headstage and directly sent to the Intan 512ch/1024ch recording controller. Singleunit offline detection was performed by thresholding the digitally filtered signal (0.8 – 5 kHz) over 5 standard deviations from the root mean square in 0.2 ms sliding windows. For each single-unit, 32 data points (1.33 ms) were sampled. A principal component analysis was implemented to extract the first three components of each spike waveform for each tetrode channel(Csicsvari et al., 1998).

Spike waveforms from individual neurons were detected using the KlustaKwik automatic clustering software(Kadir et al., 2014). Using the Klusters software(Hazan et al., 2006), single units were isolated manually by verifying the waveform shape, waveform amplitude across tetrode’s channels, temporal autocorrelation (to assess the refractory period of a single-unit) and cross-correlation (to assess a common refractory period across single-units). The stability of single-units was confirmed by examining spike features over time.

### Histology

To confirm the position of the recording sites, rats were deeply anaesthetised with urethane and lesions were made at the tip of the tetrodes using a 30 μA unipolar current for 5 – 10 s (Stimulus Isolator, World Precision Instruments). Then, rats were perfused with saline followed by 20 min. fixation with 4% paraformaldehyde, 15% (v/v) saturated picric acid and 0.05% glutaraldehyde in 0.1 M phosphate buffer. Serial coronal sections were cut at 70 or 100 μm with a vibratome (Leica). Sections containing a lesion were Nissl-stained. One rat, for which histological data could not be confirmed, was included based on: insertion coordinates, oscillatory LFP profile and similarity of neuronal activity.

### Firing rate maps and trajectory linearisation

To compute firing rate maps found in figure 1, bins of 10×10 cm were created. For each bin, the total number of spikes was divided by the rat’s occupancy (in seconds): the firing rate maps were smoothed by convolving them into two dimensions with a Gaussian low-pass filter. For the EPM, by using the geometry of the maze, the centre and the arms were defined. Trajectories were then found by demarcating the consecutive tracked positions going from the furthest point reached on the arm, to the furthest point reached on the next visited arm. To linearise this position, each two-dimensional point (x, y) was projected to the directional vector describing the arm to which that point belongs. Each projection was made by using the following equation:

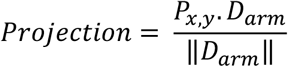

Were P_x,y_ is the position to be projected, D_arm_ is the directional vector of the corresponding arm and ||D|| denotes the norm of the vector D. Each trajectory was composed of three parts: Starting arm, centre and ending arm. For the starting and ending arm, the activity was calculated over the space by dividing the total number of spikes on each linear bin (5 cm) over the occupancy (in seconds) on that particular bin. However, due to the different possible trajectories that the animal can follow in the centre, the activity there was divided into five fixed time bins. Then, the linear firing rate maps (activity in the starting, centre and ending arm) were smoothed by convolving them with a 1-D Gaussian function. Linear firing rate maps on the ELM were calculated by dividing the space into bins (2.5 cm each) and for each bin, the corresponding spikes of each neuron were summed and divided by the occupancy (in seconds).

### Coverage index

The coverage term used is the same as the sparsity term used by Skaggs in 1996(Skaggs et al., 1996).

The formula is:

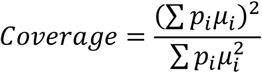

where the spatial bins of the environment are denoted by i, pi is the probability of occupying that bin and μi is the mean firing rate of the neuron in that particular bin.

### Place field similarity

The place field similarity was calculated by using the Spearman correlation between the z-scored linear firing rate map of a neuron while the animal is moving in one direction and the z-scored linear firing rate map of the same neuron while the animal is moving in the other direction.

### General Linear Model and neuronal spike activity relation to speed

We modelled the spike activity of each neuron (for different ELM configurations) by using the instantaneous running speed of the animal at each moment: *Spk_t_* = *β*_0_ + *β*_1_*S_t_* where *Spk* is the number of spikes at a given time *t* and *S* is the instantaneous running speed at the same time *t*. Residuals of the model were used as the spike-associated activity of the neurons, controlled by the speed of the animal.

### Neuronal state-space

By using the firing rate of each neuron prior to the entry to the open arm in the CO configuration, each trial was represented by a population vector built with the activity of all the co-recorded neurons during that particular day *T_n_* = {*FR*_1*n*_,*FR*_2*n*,_, …, *FR_mn_*) where FR is the firing rate, n is the total number of trials, and m is the total number of neurons.

### SVM classifier and neuronal selection

The exploration of the open arm on the ELM was divided into either proximal exploration or distal exploration using a threshold per session 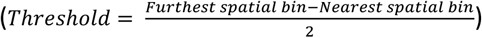. A support vector machine classifier with a linear kernel was used to determine if, based on the neuronal activity during the closed arm of the CO configuration, the extent of the exploration of a given trial (proximal or distal exploration) was possible. To do so, in a “one-leave out cross-validation” fashion, the identity of each trial was predicted by training the classifier with the activity of all the other trials. We then iteratively check the performance of the SVM for all possible combinations of neurons for each session. The neuron combination that gave the highest performance was then assumed to be optimal. In case that more than one possible combination was giving the exact same value of performance, the neurons with the higher number of appearances in the combinations (75% quartile of the repetition distribution) were selected as the set of “informative” neurons.

Due to the fact that the distal and proximal trial distributions were not even, in order to determine if a given performance value was in fact better than random, we shuffle 1000 times the trial IDs (distal or proximal) and repeated the classification. Only performance values above the 95% percentile of the shuffle distribution were considered as successful.

Two recording sessions were excluded due to either a low number of co-recorded cells (n < 3) or insufficient exploration of the maze.

### Statistical Analyses

All calculations were made in MATLAB (Mathworks, version R2015b and R2019b) and statistical analyses were performed with MATLAB and Microsoft Excel. All the statistical tests used in this manuscript were non-parametric unless stated otherwise. Raw data was visualised and visually evaluated with Neuroscope (http://neurosuite.sourceforge.net/information.html).

One-Way ANOVA with Tukey-Kramer for multiple comparison was used for the following analyses: Time spent on each of the EPM and ELM areas (Figure 1A, Figure 2E). Wilcoxon signed-rank test was used for: time spent in changing and non-changing arms (Figure 2D); spatial broadening of neuronal activity (Figure 3E). Chi-squared test was use for: Proportions of peak firing activities (Figure 3B, C and Figure S1A, B). Two-sample Kolmogorov-Smirnov test was used for comparing the PFS index distributions (Figure 4B, C and Figure S1C, D).

Analyses of CC and CO have an n = 133, while analyses of CC2 and CT have an n = 75. Both numbers correspond to the recorded neurons in the vH. Time spent measurements were done per session. EPM sessions (n = 16), ELM Sessions (n = 14). Control sessions of CC2 and CT (n = 5).

## ADDITIONAL INFORMATION

### Lead contact

Further information and requests may be directed to the Lead Contact, Thomas Klausberger.

### Data and code availability

Any original code or additional information required to reanalyse the data reported in this paper is available from the lead contact upon request.

## Acknowledgements

This work was supported by grant P 29588 of the Austrian Science Fund (T.K.), ERC starting grant 716761 (S.C.) and a Swiss National Science Foundation professorship grant (170654) (S.C.).

## Contributions

H. M-V., S.C. and T.K. contributed to experiments, data analysis and the preparation of the manuscript.

